# LOTUS-domain proteins activate Vasa to drive germ granule assembly

**DOI:** 10.64898/2026.01.08.698407

**Authors:** Ian F. Price, Chin Wang, Wen Tang

**Affiliations:** Department of Biological Chemistry and Pharmacology, The Ohio State University, Columbus, OH 43210, USA; Center for RNA Biology, The Ohio State University, Columbus, OH 43210, USA; Ohio State Biochemistry Program, The Ohio State University, Columbus, OH 43210, USA; Molecular, Cellular and Developmental Biology Program, The Ohio State University, Columbus, OH 43210, USA

**Keywords:** LOTUS-domain proteins, DEAD-box proteins, germ granules, Vasa, intermitochondrial cement

## Abstract

Germline development relies on perinuclear membraneless germ granules, yet the mechanisms underlying their assembly remain incompletely understood. Here we uncover a conserved and central role for LOTUS-domain proteins in driving germ granule assembly. In *C. elegans*, the LOTUS-domain protein EGGD-1/MIP-1 at sub-stoichiometric levels recruits Vasa protein GLH-1 to the nuclear periphery. Acting as a catalyst, EGGD-1 impacts the ATPase cycles of GLH-1 by preferentially binding to its open conformation, enhancing its RNA binding activities, and facilitating its transition to the closed conformation. GLH-1 in the closed state enriches mRNAs at the nuclear periphery, which enables the accumulation of RNA-binding proteins including PGL-1 and PGL-3. Using human cells, we demonstrate the LOTUS-domain protein TDRD5 similarly recruits DDX4, the human Vasa homolog, and stimulates the formation of intermitochondrial cement. Collectively, these findings reveal evolutionarily conserved stimulatory effect of LOTUS-domain protein in Vasa activity and provide a unified model for germ granule assembly across species.

## Introduction

Germ lines play a crucial role in transmitting genetic and epigenetic information across generations. The development and maintenance of germ lines require perinuclear germ granules, which are known as P granules in worms, nuage in flies, intermitochondrial cement and chromatoid bodies in mammals^1–4^. Biophysical studies coupled with super-resolution microscopy have revealed that germ granules exhibit behavior of biomolecular condensates formed by phase separation^5–7^. While various biomolecular condensates, including nuclear speckles and stress granules, have been identified^5^, germ granules remain among the most enigmatic biomolecular condensates due to their dynamic nature, unique perinuclear localization, and exclusive expression in the germ line. Despite intensive investigation, the molecular principles governing germ granules assembly are not fully understood.

Biochemical and genetic approaches have identified over 100 P granules proteins in *C. elegans*^3^. Among them are LOTUS-domain (<u>L</u>imkain /MARF1, <u>O</u>skar and <u>Tu</u>dor domain-containing protein<u>s</u> 5 and 7) containing proteins EGGD-1 and EGGD-2 (<u>E</u>mbryonic and <u>G</u>ermline P <u>G</u>ranule <u>D</u>etached), also referred to as MIP-1 and MIP-2 respectively in an independent study^8–12^. LOTUS domains are found in Oskar in flies, as well as TDRD5 and TDRD7 (Tudor domain-containing proteins) in mammals^13^. Despite minimum sequence homology, LOTUS domains share a common helix-turn-helix conformation^11,12,14,15^. Broadly, LOTUS domains are categorized into two subclasses: minimal LOTUS (mLOTUS) which contains a three-helix core but lacks a C-terminal extension, and extended LOTUS (eLOTUS) which contains the core as well as C-terminal alpha-helix^15^. The mLOTUS domain is thought to associate with RNAs, while the eLOTUS domain of Drosophila *Oskar* is found to interact specifically with Vasa, a DEAD-box protein^14–16^.

In *C. elegans*, LOTUS domain-containing proteins EGGD-1 and EGGD-2 are partially redundant to facilitate the perinuclear localization of P granules^8–10^. Our previous findings showed that EGGD-1 is intrinsically capable of self-assembling into perinuclear granules^8^. Through its eLOTUS domain, EGGD-1 recruits GLH-1—a *Drosophila* Vasa homolog^17–19^, to the nuclear periphery^8,9^. GLH-1 in *C. elegans*, Vasa in *Drosophila melanogaster*, and its mammalian homolog DDX4 belong to the DEAD-box RNA helicase family. Each possesses a glycine-rich IDR (Intrinsically <u>D</u>isordered <u>R</u>egion) and a helicase core^19^. The IDR potentially interacts with nuclear pore complexes and facilitates the formation of phase-separated organelles in vitro^20–22^. The helicase core is composed of one N-terminal and one C-terminal RecA domain for binding RNAs and ATP^23,24^. Loss of either GLH-1 or EGGD-1 results in the disassembly of P granules and reduced fertility^8,9,18^, yet the functional significance of the interaction between GLH-1 and EGGD-1 remains largely unclear.

This study combines in vivo reconstitution and in vitro biochemistry to dissect the interactions between major P granule proteins and RNAs. Specifically, we show that EGGD-1 at sub-stoichiometric levels recruits GLH-1 to the nuclear periphery. GLH-1 switch between open (apo) and closed (ATP/RNA-bound) conformation, with the closed conformation favoring phase separation^20,21,23,25,26^. Our findings reveal that EGGD-1 preferentially binds to GLH-1 in the open conformation. Serving as a catalyst, EGGD-1 stimulates RNA binding and ATPase activities of GLH-1 and promotes the formation of the closed GLH-1 conformation. By enhancing GLH-1 RNA binding activity, EGGD-1 enriches the core P granule proteins including PGL-1 and PGL-3 at perinuclear granules. Additionally, TDRD5, a human LOTUS domain-containing protein, recruits and promotes condensation of DDX4, a human Vasa homolog. Taken together, our findings establish the LOTUS-domain protein–Vasa axis as a central and conserved driver of germ granule assembly from worms to mammals.

## Results

### EGGD-1 eLOTUS domain binds to GLH-1 C-terminal RecA domain

The assembly of germ granules in *C. elegans* depends on EGGD-1 and GLH-1^8,9^, yet the precise nature of their interaction remains unclear. We therefore employed structure prediction and comparative structural analysis to determine the molecular basis of the EGGD-1–GLH-1 interaction. In *D. melanogaster*, Oskar serves as the scaffold for germ plasm assembly^27^. Structural studies have revealed that the eLOTUS domain of Oskar binds to the C-terminal RecA domain of Vasa (Fig. 1a)^14,15^. Unlike *Oskar* which contains only one eLOTUS domain, EGGD-1 in *C. elegans* possesses two eLOTUS domains (Supplementary Fig. 1a). Like *D. melanogaster* Vasa, *C. elegans* GLH-1 contains a helicase core comprising N-terminal and C-terminal RecA domains (Supplementary Fig. 1b).

**Figure 1.**
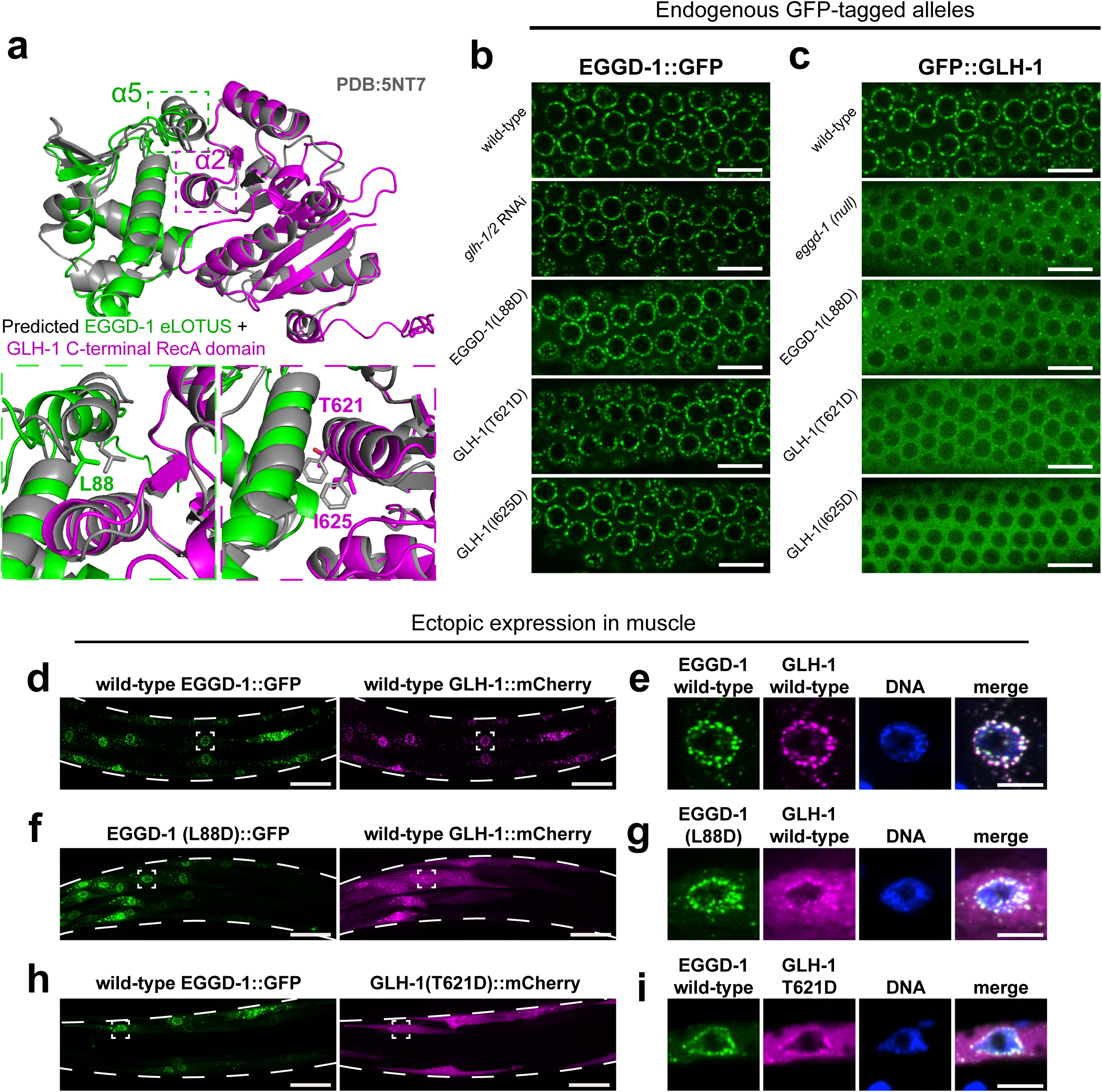
EGGD-1 eLOTUS domain interacts with GLH-1 C-terminal RecA domain. a. Structural comparison between Oskar-Vasa and EGGD-1-GLH-1. The top panel depicts the crystal structure of the interface between *D. melanogaster* Oskar eLOTUS domain with Vasa helicase C-terminal domain in gray. The predicted interface from Alphafold multimer between *C. elegans* EGGD-1 and GLH-1 is shown in green and magenta, respectively. The bottom left panel highlights a conserved interfacial residue, EGGD-1(L88), located on the α5 helix of the EGGD-1 LOTUS1 domain. The bottom right panel shows T621 and I625 in α2 helix of GLH-1 C-terminal RecA domain. b. Single-slice confocal fluorescence images showing the localization of endogenously-tagged EGGD-1::GFP around pachytene germline nuclei in wild-type as well as in the indicated mutant backgrounds and RNAi conditions. Images are representative of at least 8 worms for each background. Scale bar = 10 μm. c. Same as in (B) but depicting GFP::GLH-1. Scale bar = 10 μm. d, f, h. Maximum intensity projections of confocal z-stacks depicting body wall muscle of fixed worms co-expressing EGGD-1::GFP and GLH-1::mCherry with the indicated mutations from the muscle-specific *myo-3* promoter. Long dashed lined indicate the edges of the worms. Images are representative of at least 10 worms images for each condition. Scale bar = 20 μm. e, g, i. Single-slice confocal images of individual muscle nuclei indicated by the boxes in panels (d), (f) and (h). Scale bar = 5 μm.

Using AlphaFold prediction^28^, we identified a specific interaction between GLH-1 and the first eLOTUS domain (eLOTUS1) of EGGD-1, but not with its second eLOTUS domain (eLOTUS2). The predicted EGGD-1 eLOTUS1 and GLH-1 C-terminal RecA domain structures were further aligned onto the crystal structure of the Oskar eLOTUS–Vasa RecA complex (PDB: 5NT7)^15,28^. Consistent with previous structural modeling results^9^, the eLOTUS1 and C-terminal RecA domains form a potential interface for EGGD-1–GLH-1 interaction. In this alignment, the GLH-1 RecA domain interacts with two helices of the eLOTUS1 domain: the α2 helix which is part of the three-helix core and the α5 helix which is outside of the core (Fig. 1a)^14,15^. Conversely, EGGD-1 eLOTUS1 is contacted by the α2 helix of the GLH-1 C-terminal RecA domain (Fig. 1a).

To validate the predicted EGGD-1–GLH-1 interface, we used CRISPR/Cas9 to introduce point mutations at the putative interaction sites (Fig. 1a). Specifically, a leucine residue in the α5 helix of the EGGD-1 eLOTUS1 domain was mutated into aspartic acid (EGGD-1 L88D), while a threonine or isoleucine in the α2 helix of the GLH-1 C-terminal RecA domain was mutated to aspartic acid, referred to as GLH-1(T621D) and GLH-1(I625D) respectively (Fig. 1a,b,c). In line with previous findings^8,9^, both wild-type EGGD-1::GFP and wild-type GFP::GLH-1 granules localized to the nuclear periphery (Fig. 1b,c), with approximately 25.7% of GLH-1 observed in condensed fractions in the germ line (Supplementary Fig. 1c). EGGD-1::GFP maintained its perinuclear localization upon depletion of *glh-1* and its paralog *glh-2.* In contrast, loss of *eggd-1* led to diffusion of GFP::GLH-1 into the cytoplasm and reduced GFP::GLH-1 condensates (Fig. 1b,c). Mutations at the predicted interface of the EGGD-1–GLH-1 complex, including EGGD-1(L88D) and GLH-1(T621D) and GLH-1(I625D), did not appear to affect EGGD-1 localization (Fig. 1b). However, in all mutant backgrounds GFP::GLH-1 failed to localize to the nuclear periphery and became diffuse, with less than 1% remaining in the condensed phase (Fig. 1c and Supplementary Fig. 1c). Together, these structural and mutagenesis analyses suggest that EGGD-1, through its eLOTUS domain, recruits GLH-1 to form perinuclear granules.

To further confirm the EGGD-1–GLH-1 interaction, we employed an in vivo reconstitution system in which individual proteins were ectopically expressed under the muscle-specific *myo-3* promoter. This system allowed us to examine EGGD-1 and GLH-1 relationship in muscle cells independently of other germline proteins^8^. Using this ectopic expression system, we found that EGGD-1::GFP by itself assembled into perinuclear granules (Supplementary Fig. 1d,e), while mCherry::GLH-1 alone was diffuse throughout muscle cells (Supplementary Fig. 1f,g). Co-expression of both proteins led to the formation of EGGD-1–GLH-1 perinuclear granules (Fig. 1d,e)^8^. In contrast, EGGD-1 (L88D) mutation in the α5 helix of EGGD-1 eLOTUS1 did not robustly recruit GLH-1 (Fig. 1f,g), suggesting eLOTUS1 plays an important role in recruiting GLH-1. Similarly, GLH-1 (T621D) mutation in the GLH-1 C-terminal RecA domain did not affect EGGD-1 localization, but prevented GLH-1 from concentrating at the nuclear periphery (Fig. 1h,i). Taken together, the ectopic expression experiments demonstrate that the interaction with EGGD-1 is sufficient to recruit GLH-1 to the nuclear periphery.

### EGGD-1 at sub-stoichiometric levels recruits GLH-1 to the nuclear periphery

Since our structural model indicated a 1:1 binding ratio between EGGD-1 and GLH-1 (Fig. 1a), we expected that EGGD-1 would be expressed at levels equal to or higher than GLH-1. To investigate this, we employed three approaches to quantify the relative abundance of endogenous EGGD-1 and GLH-1 proteins. First, we examined recent proteomic data generated from isolated *C. elegans* gonad^29^. Based on four biological replicates, we found that the abundance of GLH-1 protein (Median abundance: 1.85 × 10^9^) was approximately 58-fold higher than that of EGGD-1 protein (Median abundance: 3.2 × 10^7^) (Supplementary Fig. 2a). Second, we examined strains expressing EGGD-1::GFP and GFP::GLH-1 from their respective endogenous loci, and measured GFP fluorescence by confocal microscopy. With the identical exposure time and laser intensity, GFP::GLH-1 signals were 29.3-fold stronger than EGGD-1::GFP signals within germ granules (Fig. 2a and Supplementary Fig. 2b). Finally, *3xflag* sequences were inserted to either *eggd-1* or *glh-1* genomic loci. Expression levels of EGGD-1::3xFLAG and 3xFLAG::GLH-1 were quantified through Western blotting. By diluting 3xFLAG::GLH-1 lysates (Fig. 2b) or using varying numbers of worms expressing individual FLAG-tagged proteins (Supplementary Fig. 2c), we confirmed that 3xFLAG::GLH-1 was expressed at a level at least 20-fold higher than EGGD-1::3xFLAG.

**Figure 2.**
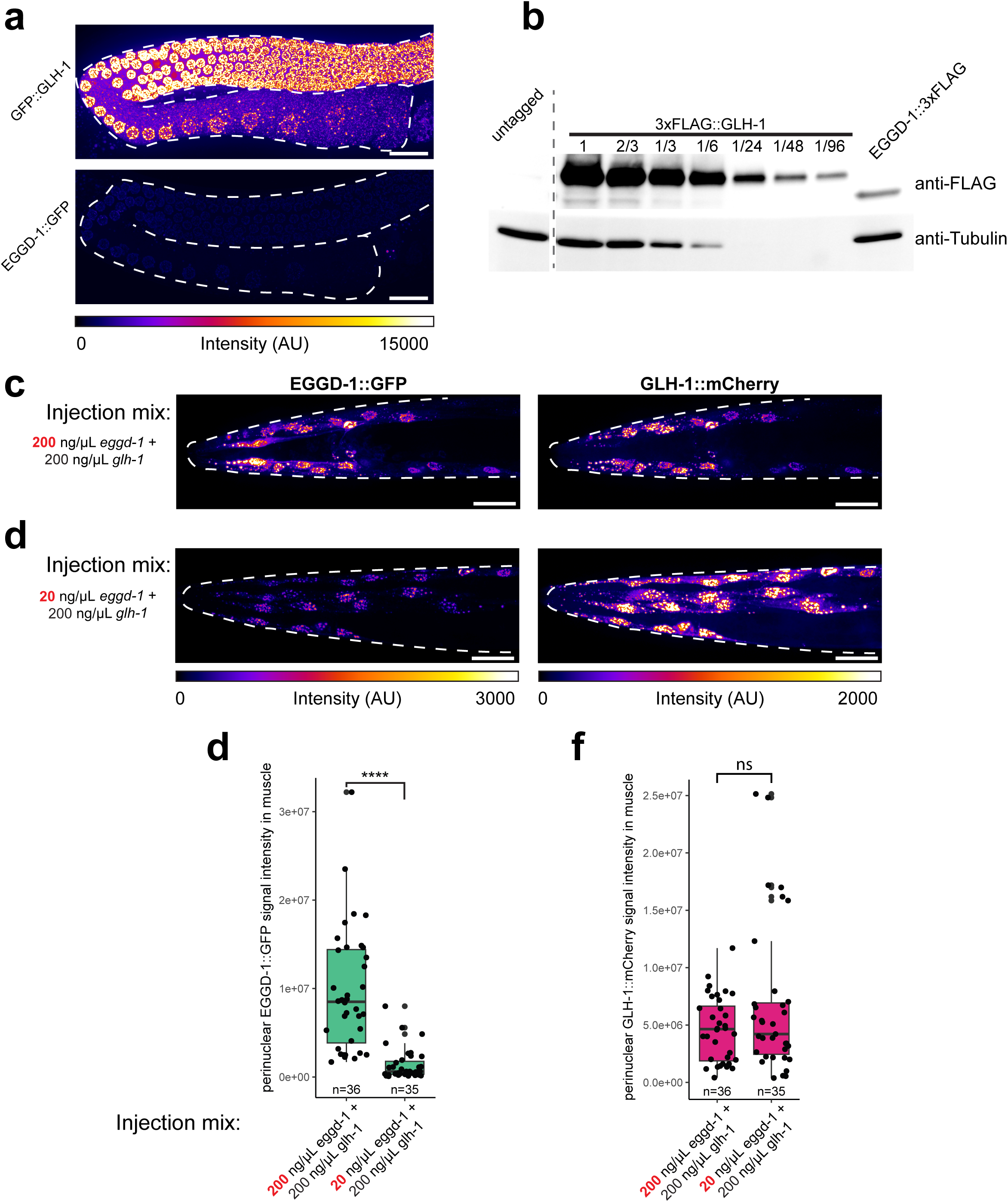
Sub-stoichiometric EGGD-1 recruits GLH-1 to the nuclear periphery. a. Maximum intensity projections of confocal z-stacks spanning the germ line showing the expression level of endogenously-tagged EGGD-1::GFP and GFP::GLH-1. The lookup table bar (bottom) indicates the intensity values for the colors depicted in arbitrary units (AU). Images are representative of at least 10 worms. Scale bar = 20 μm. b. Western blot showing the abundance of endogenously tagged EGGD-1::3xFLAG and 3xFLAG::GLH-1 in lysate from gravid adult worms. Fractions above GLH-1 bands indicate the relative dilution factor of GLH-1 lysate. Blot is representative of two independent experiments. c. Maximum intensity projection of confocal z-stacks depicting the head of worms transgenically co-expressing EGGD-1::GFP and GLH-1::mCherry from the muscle-specific *myo-3* promoter. The strain was generated using an injection mix containing 200ng/μL EGGD-1 with 200 ng/μL GLH-1 plasmid. Images are representative of two independent transmission lines with at least 8 worms imaged for each. Scale bar = 20 μm. d. Same as in (c) but the strains were generated with a mix of 20ng/μL EGGD-1 with 200 ng/μL GLH-1 plasmid. Images are representative of two independent transmission lines with at least 8 worms imaged for each. Scale bar = 20 μm. e. Quantification of EGGD-1::GFP signal around individual muscle nuclei in transgenic strains generated with a 200 ng/μL EGGD-1+200 ng/μL GLH-1 plasmid mix or a 20 ng/μL EGGD-1+200 ng/μL GLH-1 plasmid mix. n indicates number of nuclei quantified. Student’s t-test, p-value ≤ 0.00005. f. Same as in (e), but quantifying perinuclear GLH-1::mCherry signal around the same nuclei. n indicates number of nuclei quantified. Student’s t-test, p-value ns ˃ 0.05.

This substantial difference in protein abundance led us to hypothesize that EGGD-1, even at sub-stoichiometric levels, recruits GLH-1 to the nuclear periphery. We tested this hypothesis using the ectopic expression experiment, where the relative stoichiometry of EGGD-1 and GLH-1 could be adjusted by varying the plasmid concentration in the injection mix for transgenic animals. With a 1:1 plasmid ratio of *myo3-promoter::eggd-1::gfp* and *myo3-promoter::glh-1::mCherry* (200 ng/µL each), we observed co-localization of EGGD-1 and GLH-1 at the nuclear periphery (Fig. 2c). When ratio was adjusted to 1:10 (20 ng/μL *myo3-promoter::eggd-1::gfp* and 200 ng/μL *myo3-promoter::glh-1::mCherry* plasmid in the injection mix), less EGGD-1::GFP was expressed (Fig. 2c,e). However, GLH-1::mCherry was still robustly recruited to the nuclear periphery (Fig. 2d,f). These findings strongly suggest that sub-stoichiometric EGGD-1 can effectively recruit GLH-1 to the nuclear periphery.

### EGGD-1 may preferentially interact with GLH-1 in its open conformation

We next aimed to uncover how sub-stoichiometric levels of EGGD-1 interact with GLH-1. Both *C. elegans* GLH-1 and *Drosophila* Vasa are members of the DEAD-box protein (DDX) family, which is characterized by the conserved Asp-Glu-Ala-Asp (DEAD) motif and two RecA domains (Supplementary Fig. 1b)^18,24,30^. DEAD-box proteins including GLH-1 and Vasa undergo a dynamic transition between closed and open conformations. In the closed conformation, the two RecA domains bind to RNAs and ATP (Fig. 3a). In this ATP/RNA-bound state, DEAD-box proteins clamp on RNA, leading to the formation of higher-order biomolecular condensates^24^. Upon ATP hydrolysis, DEAD-box proteins shift to the open conformation, where the RecA domains become separated and inorganic phosphate, ADP and RNA are released (Fig. 3a). This transition facilitates the repositioning of RNA strands, allowing DEAD-box proteins to locally unwind RNA duplexes or remodel RNA-protein complexes^24,31^.

**Figure 3.**
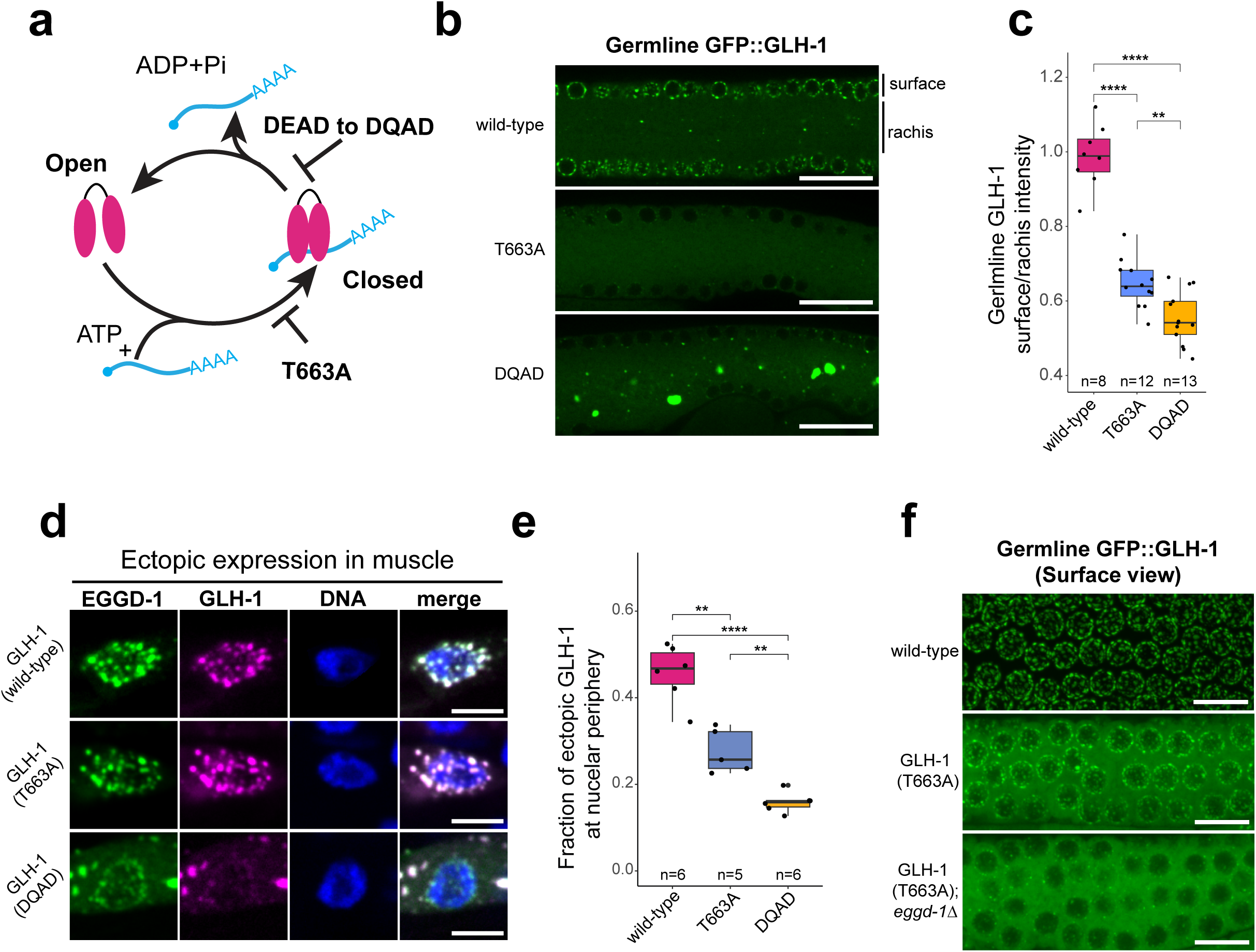
EGGD-1 recruits open-conformation GLH-1 to the nuclear periphery. a. Schematic of the GLH-1/DEAD box protein ATPase cycle. open conformation GLH-1 associates with RNA in an ATP-dependent manner and transitions to a closed conformation. ATP hydrolysis destabilizes the complex and drives disassembly, releasing ADP, inorganic phosphate, and RNA. The mutation T663A on motif V of the helicase domain blocks the association of GLH-1 with RNA. Mutation of the DEAD motif to DQAD stabilizes the ATP hydrolysis products in the helicase domain, causing prolonged association with RNA and constitutive closed conformation GLH-1. b. Single confocal slices depicting cross-sections in germ lines of live worms expressing endogenously-tagged GFP::GLH-1 (wild-type, T663A, or DQAD). Images are representative of at least 8 worms. Scale bar = 20 μm. c. Quantification of the signal ratio of the average GFP::GLH-1 intensity at the surface over the rachis within a germline cross section. Note that the nuclei are located at the edge, so the reduction of surface-to-rachis ratio indicates a relative loss of GLH-1 localized to the nuclear periphery. n = number of germ lines quantified. Student’s t-test, **p-value ≤ 0.005, ***≤0.0005, ****≤0.00005. d. Single confocal slices depicting muscle nuclei in fixed worms ectopically expressing EGGD-1::GFP and GLH-1::mCherry (wild-type, T663A or DQAD). Images are representative of at least 8 worms. Scale bar = 5 μm. e. Quantification of the total fraction GLH-1::mCherry signal localized to the periphery of EGGD-1::GFP positive nuclei in the muscle of transgenic EGGD-1::GFP; GLH-1::mCherry worms. n = number of worms quantified. Student’s t-test, ** p-value ≤ 0.005, ***≤0.0005. f. Maximum intensity projections of z-stacks encompassing the top row of germline nuclei in live worms expressing endogenously tagged GFP::GLH-1 in the indicated mutant backgrounds. Images are representative of at least 8 worms. Scale bar = 10 μm.

Previous studies have shown that GLH-1 conformational states can be manipulated by specific amino acid substitutions^20,21,26^. For example, a conserved threonine in the C-terminal RecA domain directly contacts with the RNA backbone^23^. When mutated to alanine (T663A), the RNA binding capacity of GLH-1 is compromised, causing it to remain in the open conformation (Fig. 3a)^21,23^. In this scenario, GFP::GLH-1(T663A) diffused into the cytoplasm (Fig. 3b). Conversely, a DQAD mutation in the DEAD motif inhibits the release of RNA and ATP hydrolysis products, which locks GLH-1 in the closed conformation and leads to the formation of large GLH-1 cytoplasmic aggregates (Fig. 3a,b)^20,21,26^. *C. elegans* germ nuclei are situated on the surface of gonad and share a central cytoplasmic core known as rachis^32^. To evaluate perinuclear localization, we quantified GFP::GLH-1 signals at the germline surface and rachis (Fig. 3b,c). This surface-to-rachis ratio reflects the enrichment of GFP::GLH-1 at the nuclear periphery. While this ratio was approximately 0.988 in wild-type, it was reduced to 0.646 in GFP::GLH-1(T663A) mutants and further reduced to 0.554 in GFP::GLH-1(DQAD) mutants (Fig. 3c). These findings indicate that the ATPase cycle of GLH-1 is crucial for its perinuclear localization.

Since GLH-1 in the open conformation showed a less severe recruitment defect than the closed conformation (Fig. 3b,c), we hypothesized that EGGD-1 may preferentially associate with the open conformation of GLH-1. We employed the ectopic expression system to directly test this hypothesis. T663A and DQAD substitutions were introduced to *myo3-promoter::glh-1::mCherry* plasmid. When EGGD-1 was co-expressed with GLH-1 mutants, it recruited GLH-1(T663A) to the nuclear periphery (Fig. 3d,e and Supplementary Fig. 3). However, EGGD-1 failed to effectively recruit GLH-1(DQAD) (Fig. 3d,e and Supplementary Fig. 3). To further validate the interaction between open GLH-1 and EGGD-1, we assessed the localization of endogenous GFP::GLH-1(wild-type), GFP::GLH-1(T663A), and GFP::GLH-1(T663A) in the *eggd-1* mutant background. While GLH-1(T663A) was mostly diffuse, a small fraction remained perinuclear in wild-type animals (Fig. 3b,c,f). However, in the absence of EGGD-1, perinuclear GLH-1(T663A) was further reduced (Fig. 3f), suggesting EGGD-1, at sub-stoichiometric levels, can recruit open GLH-1 to the nuclear periphery. Taken together, our genetic and microscopy data indicate that EGGD-1 may preferentially bind to GLH-1 in its open conformation.

### EGGD-1 stimulates GLH-1 ATPase and RNA binding activities and promotes its transition to the closed conformation

The preferential binding of EGGD-1 to the open conformation of GLH-1 led us to hypothesize that EGGD-1 modulates GLH-1 ATPase and/or RNA binding activities. We sought to develop in vitro assays to test this hypothesis. Due to the challenges in purifying full-length GLH-1 and EGGD-1, we purified a truncated recombinant GLH-1 with intact RecA domains (Amino Acid 279-763), wild-type eLOTUS1, and eLOTUS1(L88D)—a mutant that does not robustly interact with GLH-1 (Fig. 1f, 1g and Supplementary Fig. 4).

First, we employed a malachite green assay to assess GLH-1 ATPase activity in vitro (Fig. 4a). In this assay, the release of inorganic phosphate (Pi) from ATP forms a phosphomolybdenum complex that interacts with malachite green, which causes the dye to absorb light at wavelengths around 600nm (Fig. 4a)^33^. In a time-course experiment, ATP hydrolysis was detected with GLH-1 protein, but not with the wild-type eLOTUS1 domain (Fig. 4b). When wild-type eLOTUS1 was incubated with recombinant GLH-1, ATP hydrolysis was increased substantially, while the addition of eLOTUS1(L88D) did not produce this effect (Fig. 4b). These findings indicate that the eLOTUS1 domain of EGGD-1 stimulates the ATPase activity of GLH-1.

**Figure 4.**
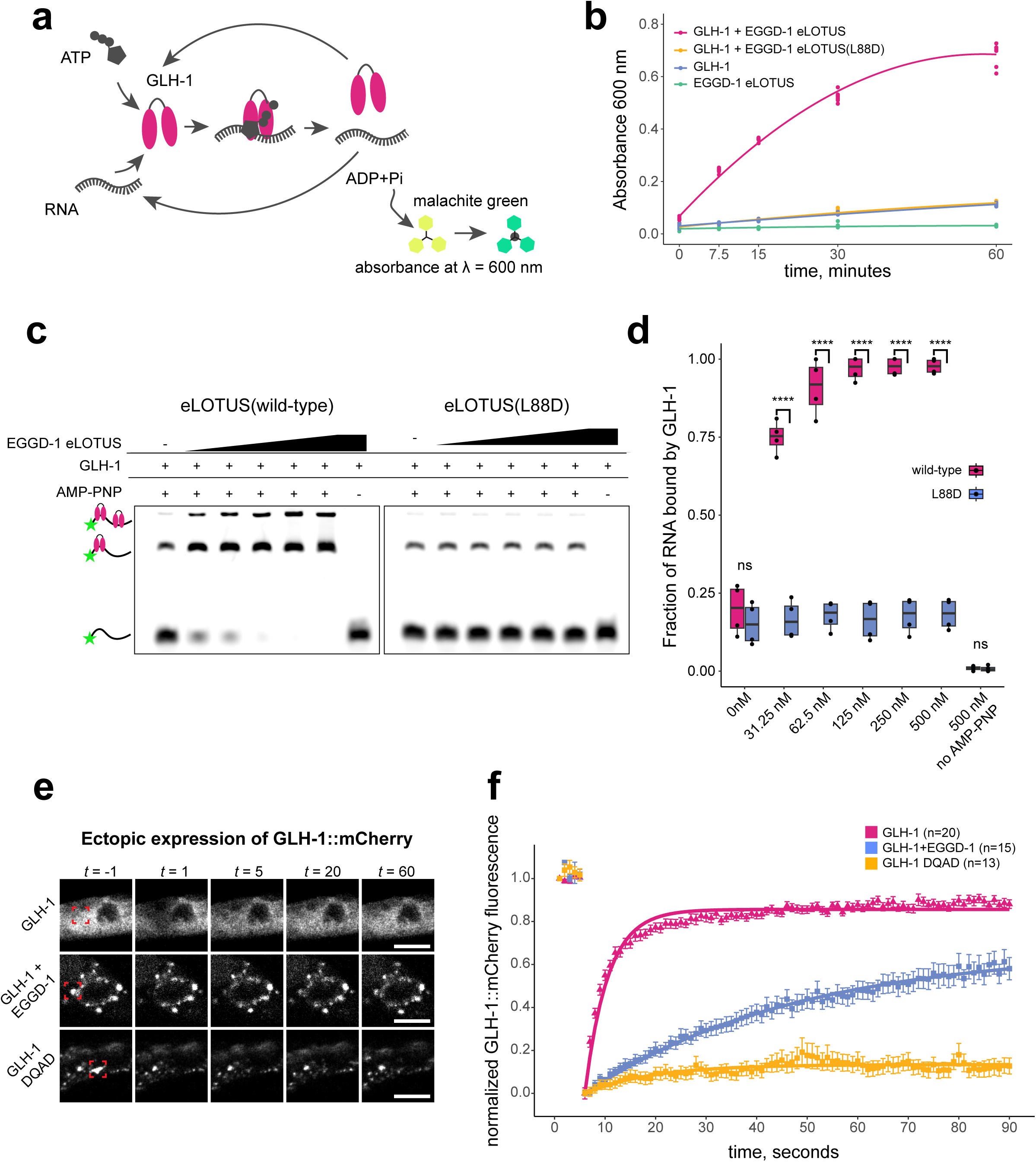
Interaction with EGGD-1 promotes GLH-1 binding to RNA GLH-1 retention in condensed foci. a. Schematic for assessing GLH-1 ATPase activity through a malachite green assay. After each ATPase cycle, GLH-1 releases inorganic phosphate which reacts with the malachite green reagent to cause absorbance at wavelengths around 600nm. b. Absorbance data from a malachite green assay to measure the production of inorganic phosphate, as indicated by absorbance at 600nm, over time in the presence of the indicated recombinant proteins. Data are generated from 4 independent reactions with two technical replicates each. c. Electrophoretic mobility shift assay examining the binding of GLH-1 to a 30 nt poly-U RNA substrate in the presence of wild-type or mutant EGGD-1 eLOTUS domain and a non-hydrolysable ATP analog, AMP-PNP. Gels are representative of 4 independent experiments. d. Quantification of the fraction of RNA bound by GLH-1 from the EMSA assay shown in panel (c). n = 4 independent experiments. Student’s t-test, **** p-value ≤ 0.00005. e. Representative examples of fluorescence recovery after photobleaching (FRAP) experiments showing the recovery of transgenic GLH-1::mCherry fluorescence signal in muscle cells expressing wild-type GLH-1, wild-type GLH-1 with EGGD-1, and DQAD mutant GLH-1 at the indicated time points. Photobleached regions are outlined by the red dashed box. Images are representative of at least 13 bleached ROIs across ≥ 5 separate worms. Scale bar = 5μm. f. Plot of normalized fluorescence recovery after photobleaching in transgenic muscle cells as shown in (e). Data points are mean recovery at the indicated times ± standard error. n = number of GLH-1::mCherry foci bleached.

Next, we examined the RNA binding activity of GLH-1 using an electrophoretic mobility shift assay (EMSA). In this assay, a FAM labeled 30-nt poly-U RNA was incubated with recombinant GLH-1 and eLOTUS proteins. Because DEAD-box proteins transiently interact with RNA^18,24,30^, we included AMP-PNP, a non-hydrolyzable ATP analog, to lock GLH-1 onto the RNA substrate. As expected, GLH-1-RNA complexes were observed only in the presence of AMP-PNP (Fig. 4c,d). Wild-type eLOTUS1 stimulated the formation of GLH-1-RNA complexes in a concentration dependent manner (Fig. 4c,d). In contrast, eLOTUS1(L88D) mutant did not promote complex formation (Fig. 4c,d), indicating that the eLOTUS domain of EGGD-1 enhances the RNA binding activity of GLH-1.

Previous studies have shown that GLH-1 in the constitutively closed conformation forms irregular and immobile foci^20,21,26^. Since eLOTUS facilitates GLH-1 ATPase and RNA binding activities in vitro (Fig. 4c,d), we hypothesized that EGGD-1 promotes the formation of closed GLH-1 in vivo, thus making GLH-1 foci less mobile. We employed fluorescence recovery after photobleaching (FRAP) to determine the mobility of GLH-1 in the ectopic expression system. In muscle cells expressing mCherry::GLH-1 alone, fluorescence recovery occurred within 20 seconds post-bleaching (Fig. 4e,f). In contrast, the GLH-1(DQAD) mutant, constitutively in the closed state, showed minimal recovery after photobleaching (Fig. 4e,f). When GLH-1 and EGGD-1 were co-expressed, GLH-1 recovered at a rate slower than GLH-1 alone, suggesting EGGD-1 reduces GLH-1 mobility, possibly by inducing closed GLH-1 (Fig. 4e,f). Taken together, our data support a model that EGGD-1 binds to GLH-1 in the open conformation and facilitates its transition to an ATP/RNA-bound closed conformation, the latter of which is less mobile.

### EGGD-1 and GLH-1 enrich mRNAs in the perinuclear granules

Because the eLOTUS domain enhances GLH-1 RNA binding activities in vitro (Fig. 4c,d), we hypothesized that such enhanced RNA-binding activity is required for enriching RNAs in germ granules. The majority of mRNAs in *C. elegans* are trans-spliced to one of two spliced leaders, with over 50% spliced to spliced leader 1 (SL1)^34^. This characteristic enabled us to assess the bulk mRNA distribution using Fluorescent in situ Hybridization (FISH) with a Cy5-labeled probe against SL1 (Fig. 5a)^35,36^.

**Figure 5:**
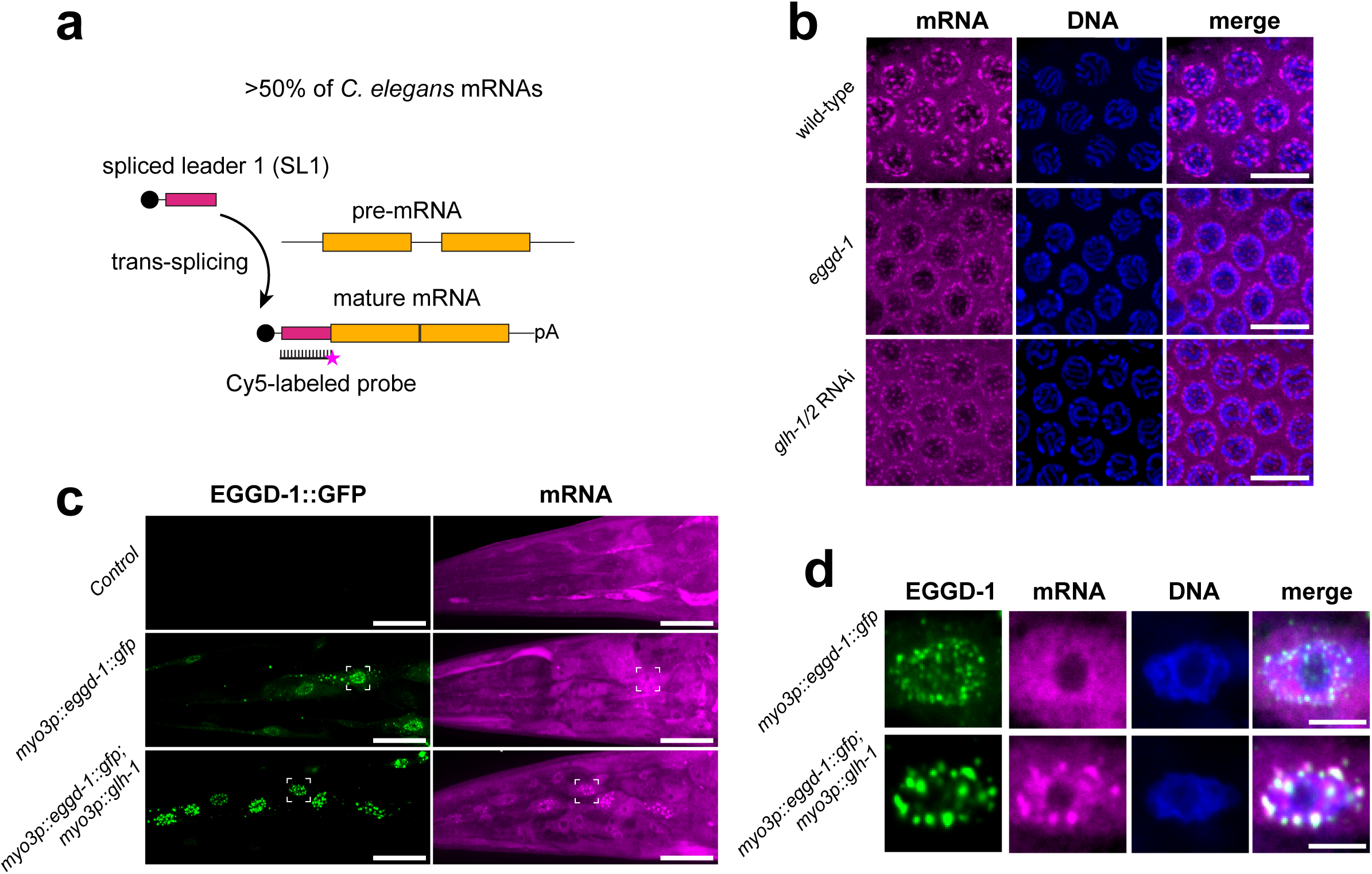
EGGD-1 and GLH-1 function to enrich mRNA at the nuclear periphery. a. Schematic showing the strategy for bulk mRNA visualization by fluorescence in-situ hybridization. Approximately 50% of *C. elegans* mRNAs are trans-spliced at the 5’ end with the 21-23 nt long spliced leader 1 RNA. A fluorescently labeled probe targeting the SL1 consensus sequence is used to visualize trans-spliced mRNAs. b. Maximum intensity projections of confocal z-stacks encompassing the top row of nuclei from gonads hybridized with a Cy5-labeled SL1 probe to visualize native SL1 foci in the indicated mutant backgrounds. Images are representative of at least 10 germlines from 3 independent experiments (≥30 total). Scale bar = 10μm. c. Maximum intensity projection of confocal z-stacks encompassing the heads of worms expressing the indicated transgenes hybridized with a Cy5-labeled SL1 probe. Images are representative of at least 10 worms from 2 independent experiments (≥20 total). Scale bar = 20μm. d. Single slice confocal image examples of nuclei indicated by the dashed boxes in (c). Scale bar = 5μm.

In wild-type germ lines, we observed that mRNAs were modestly enriched at the nuclear periphery (Fig. 5b). However, loss of either *eggd-1* or *glh-1* resulted in a reduction of mRNA signals (Fig. 5b), indicating that both EGGD-1 and GLH-1 play an important role in enriching mRNAs at the nuclear periphery. To determine whether GLH-1 activities are sufficient for mRNA recruitment, we conducted SL1 FISH in the ectopic expression system. In control animals, mRNAs were distributed throughout the muscle cells (Fig. 5c). EGGD-1 by itself formed perinuclear granules, but did not appear to enrich mRNAs (Fig. 5c,d). Notably, co-expression of EGGD-1 and GLH-1 resulted in robust mRNA enrichment in perinuclear granules (Fig. 5c,d). We conclude that EGGD-1 and GLH-1 together are necessary and sufficient for mRNA enrichment at the nuclear periphery.

### EGGD-1 activates GLH-1 RNA binding activity to recruit PGL proteins to germ granules

PGL (<u>P</u>-<u>G</u>ranu<u>L</u>e abnormality) proteins, including PGL-1 and PGL-3, are core components of *C. elegans* germ granules^37,38^. Both proteins contain RNA binding domains. Importantly, mRNA, but not ribosomal RNA, stimulates phase separation of PGL proteins in vitro^39,40^. Since EGGD-1 and GLH-1 effectively recruit mRNA to the nuclear periphery (Fig. 5), we investigated whether high concentrations of perinuclear mRNA could facilitate granule formation of PGL-1 and PGL-3 at the nuclear membrane.

First, we employed the ectopic expression system to interrogate the interplay among EGGD-1, GLH-1, PGL-1, and PGL-3. In muscle cells co-expressing EGGD-1::mCardinal and PGL-1::GFP, EGGD-1 was largely perinuclear. PGL-1 formed cytoplasmic foci, but did not colocalized with EGGD-1 (Fig. 6a,b). Upon the addition of GLH-1, PGL-1 protein was mixed with EGGD-1 and enriched at the nuclear periphery (Fig. 6a,b). The colocalization was quantified as Pearson correlation coefficient with 1 being a perfect correlation, 0 being no correlation, and –1 being perfect anti-correlation. The quantification revealed that that PGL-1 colocalized with EGGD-1 only in the presence of GLH-1 (Fig. 6a,b and Supplementary Fig. 5a). Similar results were observed with PGL-3, which formed smaller granules but was also recruited to the nuclear periphery and colocalized with EGGD-1 when GLH-1 was present (Fig. 6c,d and Supplementary Fig. 5b).

**Figure 6:**
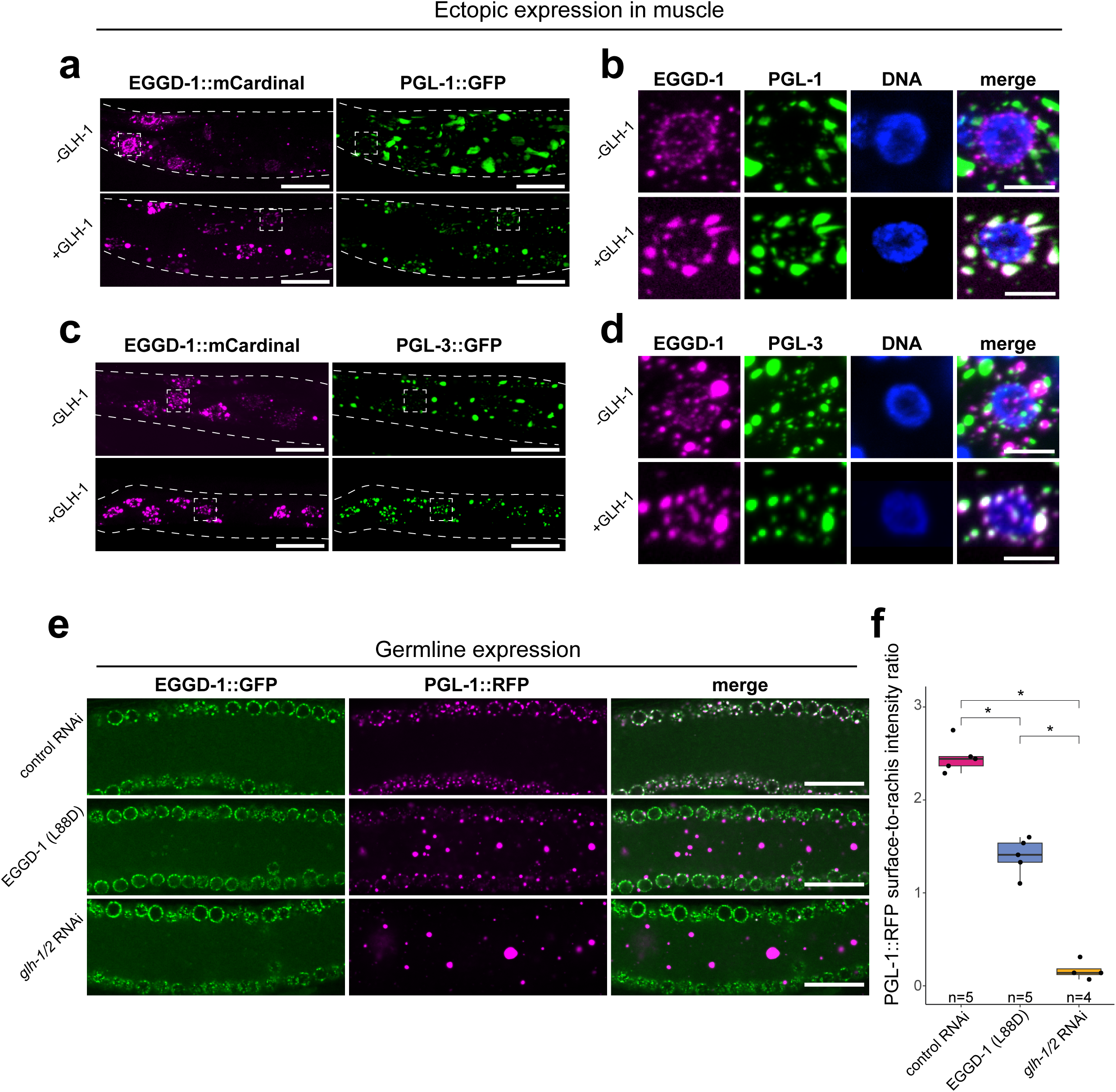
EGGD-1 and GLH-1 are necessary and sufficient to drive localization of PGL proteins. a. Body wall muscle of worms co-expressing fluorescently tagged EGGD-1 and PGL-1 in the absence or presence of GLH-1. Dashed lines outline the image worm. Images are representative of at least 10 worms from 2 independent transgenic transmission lines (≥20 total). Scale bar = 20 μm. b. Single confocal slices of specific examples of nuclei indicated by the dashed boxes in (a). Scale bar = 5 μm. c. Body wall muscle of worms co-expressing fluorescently tagged EGGD-1 and PGL-3 in the absence or presence of GLH-1. Dashed lines outline the image worm. Images are representative of at least 10 worms from 2 independent transgenic transmission lines (≥20 total). Scale bar = 20 μm. d. Single confocal slices of specific examples of nuclei indicated by the dashed boxes in (c). Scale bar = 5 μm. e. Confocal cross sections from worms expressing endogenously tagged EGGD-1::GFP and PGL-1::RFP in the indicated mutant or RNAi conditions. Images are representative of at least 4 worms. f. Plot showing surface-to-rachis intensity ratio PGL-1::RFP in the indicated mutant and RNAi conditions. n indicates number of germ lines quantified. Student’s t-test, * p-value < 0.05.

Next, we examined the distribution of germ granule proteins in animals expressing EGGD-1::GFP and PGL-1::RFP from their endogenous genomic loci. We assessed PGL-1::RFP fluorescence at both the surface and the rachis of germ line. In wild-type animals, EGGD-1::GFP and PGL-1::RFP were associated with the periphery of germ nuclei (Fig. 6e,f). When the EGGD-1(L88D) mutation was introduced, or when *glh-1* and *glh-2* were depleted, EGGD-1::GFP remained perinuclear (Fig. 6e and Supplementary Fig. 5c). In these two mutant backgrounds, the distribution of PGL-1::RFP was changed, with fewer perinuclear foci and more foci at the rachis (Fig. 6e,f). The surface-to-rachis ratio for PGL-1::RFP decreased from 2.36 in wild-type to 1.36 in EGGD-1(L88D) and 0.16 in *glh-1/2* depleted animals (Fig. 6f). Despite the presence of a few perinuclear PGL-1 foci in mutant animals, these foci were no longer colocalized with EGGD-1 (Supplementary Fig. 5d), as indicated by a Pearson correlation coefficient of –0.299 in EGGD-1(L88D) mutants and – 0.241 in *glh-1/2-*depleted animals (Supplementary Fig. 5e). Overall, our genetic and microscopy data suggest that EGGD-1, likely through its enhancement of GLH-1 mRNA-binding capability, plays a crucial role in enriching PGL proteins at perinuclear germ granules.

### TDRD5 and DDX4 form granules resembling intermitochondrial cement

Vasa proteins are conserved across the animal kingdom^19^. All Vasa proteins, including GLH-1 in *C. elegans* and DDX4 in human, contains N-terminal and C-terminal RecA domains (Supplementary Fig. 1b). AlphaFold-based protein predictions reveal that GLH-1 and DDX4 adopt highly similar structures, with a Root Mean Square Deviation (RMSD) of 0.851 (Supplementary Fig. 6a). While *C. elegans* EGGD-1 lacks a clear mammalian homolog, mammalian TDRD5 and TDRD7 (Tudor domain-containing proteins) possess the eLOTUS domain (Supplementary Fig. 1a)^13^. Notably, TDRD5 is specifically expressed in the testes, where it is highly enriched in chromatoid bodies and intermitochondrial cement (IMC)^41^. Domain analysis identified one Tudor domain, one eLOTUS and two mLOTUS domains within TDRD5 (Supplementary Fig. 1a)^13^. Although only 18.2% of the eLOTUS domain of TDRD5 is identical to that of EGGD-1 at the amino acid level (Supplementary Fig. 6b), structural predictions from AlphaFold suggest that both eLOTUS domains share a similar configuration, including an extended α5 helix (Supplementary Fig. 6c)^28^. These in silico analyses led us to hypothesize that TDRD5 interacts with DDX4 in humans.

To test this hypothesis, we ectopically expressed TDRD5 in the U2OS cell, an osteosarcoma cell line. When expressed by itself, TDRD5 intrinsically associates with mitochondria (Fig. 7a,b). Specifically, it appeared to be highly enriched on the surface of mitochondria (Fig. 7a,b,c), a pattern similar to the expression of endogenous TDRD5 in the testes^41^. In contrast, when expressed alone, DDX4 remained diffusely distributed throughout the cytoplasm (Fig. 7d). When both proteins are co-expressed, DDX4 was recruited by TDRD5 and formed IMC-like structures in U2OS cells (Fig. 7e). Next, we conducted structure modeling and identified an isoleucine residue at the α5 helix of TDRD5 eLOTUS with the potential to interact with RecA domain (Supplementary Fig. 6b,c). When we mutated this residue to aspartic acid (I87D), TDRD5 lost its ability to recruit DDX4 (Fig. 7f). Finally, If TDRD5 stimulates RNA binding of DDX4 and facilitate its transition into the closed conformation, we would expect DDX4 to be less mobile in the presence of wild-type TDRD5. To test this, we conducted a FRAP experiment on DDX4 when it was co-expressed with either wild-type TDRD5 or TDRD5 (I87D). Indeed, DDX4 recovered more slowly when co-expressed with wild-type TDRD5 compared to the I87D mutant (Fig. 7g,h). Collectively, these findings suggest that TDRD5, through its eLOTUS domain, recruits DDX4 and promotes the formation of IMC-like structures in U2OS cells.

**Figure 7:**
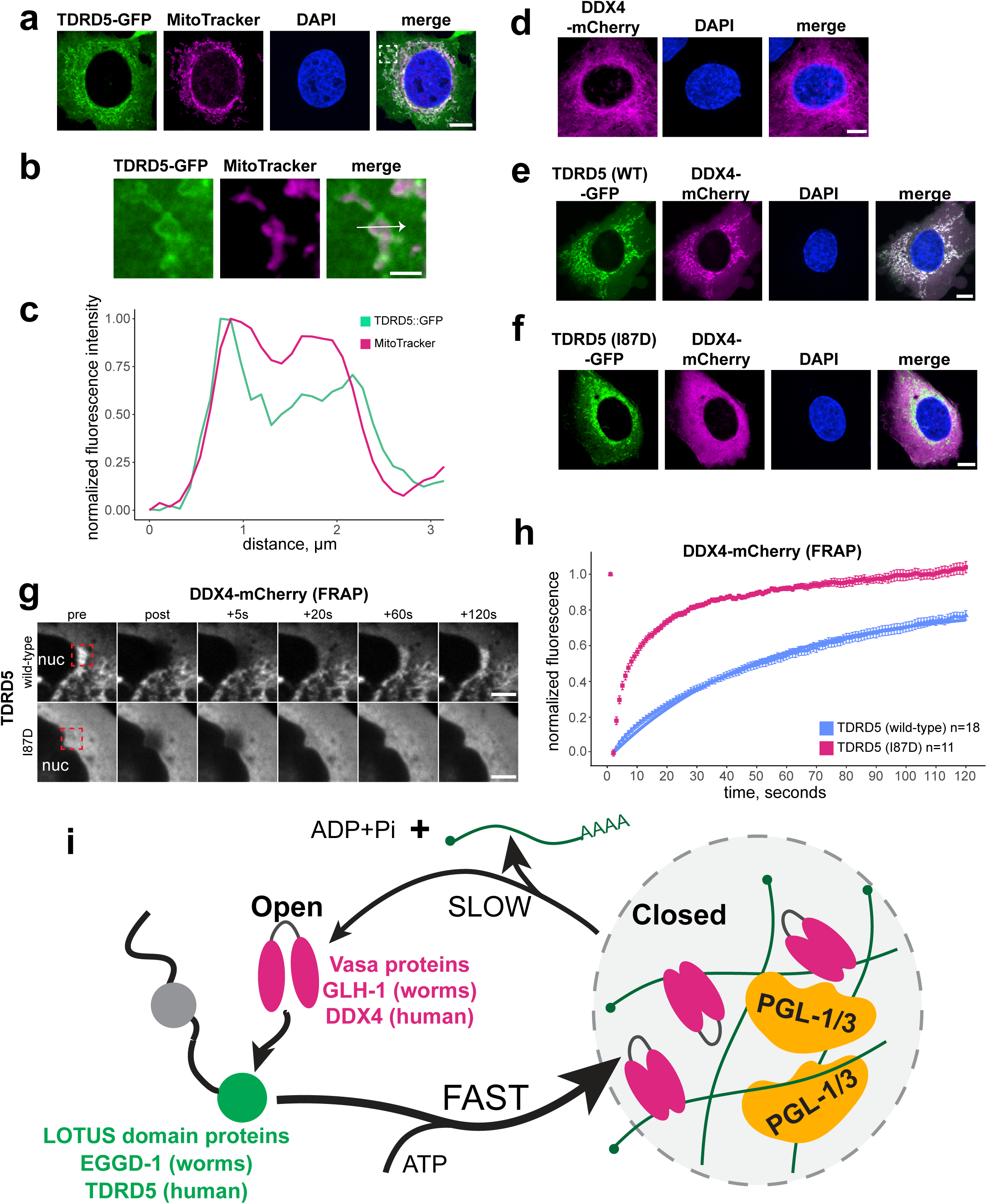
TDRD5 and DDX4 form granules resembling intermitochondrial cement. a. Representative single slice confocal image of fixed U2OS cells transfected with plasmid to express TDRD5-GFP and stained with MitoTracker Red. Images are representative of 10 TDRD5-GFP positive cells over one experiment. Scale bar = 10 μm. b. Zoomed-in view of the region within the dashed white box shown in the merged image in (a). Scale bar = 2 μm. c. Line scan plot showing the normalized fluorescence intensity of TDRD5-GFP and MitoTracker red along the line shown in the merged image in (b). d. Representative single slice confocal image of fixed U2OS cells transfected with plasmid to express DDX4-mCherry. Images are representative of at least 10 cells from two independent transfection experiments (≥20 cells). Scale bar = 10 μm. e. Representative single slice confocal image of fixed U2OS cells transfected with plasmids to co-express DDX4-mCherry with TDRD5(wild-type)-GFP. Images are representative of at least 10 cells from two independent transfection experiments (≥20 cells). Scale bar = 10 μm. f. Same as in (e) but expressing DDX4-mCherry and TDRD5(I87D)-GFP. Images are representative of at least 10 cells from one transfection experiment. Scale bar = 10 μm. g. Representative DDX4-mCherry FRAP experiments in live U2OS cells co-expressing DDX4-mCherry with wild-type or I87D mutant TDRD5-GFP. Bleached ROIs are indicated by the dashed red box at the pre-bleach timepoint. Scale bar = 5 μm. h. Plot of the fluorescence recovery curve of DDX4-mCherry after photobleaching over time. Data points are mean recovery at the indicated time. n indicates the number of ROIs bleached. i. Model showing that eLOTUS domain proteins including EGGD-1 and TDRD5 associate with Vasa proteins and stimulate their RNA-binding activity.

## Discussion

Germ cells development relies on membraneless germ granules, which maintain genome integrity and regulate gene expression^1–4^. Germ granules contain diverse RNAs and proteins. The dynamic interplay between RNA and protein components allows germ granules to form phase-separated structures. Despite progress in identifying germ granule components, outstanding questions remain: What are the mechanisms by which RNAs and proteins are recruited to germ granules? How do these components interact to achieve germ granule assembly? Are these protein-protein and protein-RNA interactions conserved across species or are they species-specific? This study provides some mechanistic insights into these questions.

Based on this work and previous studies in *D. melanogaster*^8,9,14,15^, we propose a unified model for germ granules formation (Fig. 7i): At the core of germ granule assembly, eLOTUS-domain proteins, such as EGGD-1 in *C. elegans*, Oskar in *D. melanogaster*, and TDRD5 in *H. sapiens*, serve as nucleators and/or catalysts. They recruit Vasa proteins to germ granules and promote their ATPase cycles. We show that *C. elegans* EGGD-1 preferentially binds to GLH-1 in its open conformation, possibly induces a conformation change, and facilitates its association with ATP and RNAs. ATP/RNA-bound GLH-1 transitions into a closed conformation, and therefore diffuses more slowly than GLH-1 in its open state. The net outcome is the accumulation of GLH-1 at the nuclear periphery. Importantly, the accumulation of RNAs at the nuclear periphery recruits downstream RNA-binding factors such as PGL-1 and PGL-3 proteins (Fig. 7i).

The eLOTUS domains exhibit remarkable sequence diversity, yet they share a conserved helix-turn-helix structure with an extended C-terminal alpha-helix^15^. This conserved structural feature is likely critical for their interaction with Vasa proteins across species. The sequence diversity within eLOTUS domains, on the other hand, may contribute to species-specific variations in binding affinities and hence Vasa activities. Cross-species experiments examining the interaction between eLOTUS domains and Vasa homologs may provide more insights into how these conserved functions are tailored across species.

We found the eLOTUS domain preferentially interacts with *C. elegans* Vasa in its open conformation (Fig. 3), and stimulates Vasa ATPase and RNA-binding activities (Fig. 4), potentially by inducing its conformational rearrangements. Future investigation, including structural and single-molecule studies, is required to fully understand how eLOTUS domains influence GLH-1 conformational dynamics. Notably, Modulation of DEAD-box protein activity through interactions with co-factors has been observed in other biological contexts. For example, eukaryotic translation initiation requires the eIF4F complex, which is composed of the DEAD-box protein eIF4A, the cap-binding protein eIF4E, and the scaffold protein eIF4G^42^. Analogous to the interaction between EGGD-1 and GLH-1, the scaffold protein eIF4G binds to the C-terminal RecA domain of eIF4A and stimulates its RNA binding activity^43^.

Vasa proteins belong to the DEAD-box RNA helicase family^19^. Historically DEAD-box proteins have been studied for their ATPase and RNA unwinding activities^24,31^. However, recent findings reveal that some DEAD-box proteins are non-processive and act through single-turnover reactions on RNA duplexes^44,45^. This emerging view highlights an essential role for DEAD-box proteins as RNA clamps that recruit other proteins. ^44,45^ For example, the DEAD-box protein eIF4AIII clamps RNAs within the exon junction complex (EJC), where it locks EJC onto RNA targets^46^. This study underscores the impact of the RNA-binding activity of GLH-1 in germ granule assembly. GLH-1, along with EGGD-1, is necessary and sufficient for enriching mRNAs within germ granules (Fig. 5). The enrichment of RNAs at the nuclear periphery is critical for recruiting PGL proteins (Fig. 6). These findings support a model that in the presence of EGGD-1, the primary function of GLH-1 may not be RNA unwinding, but rather clamping onto mRNA in its closed conformation. The helicase activity, one the other hand, may play a role in fine-tuning RNA-RNA and RNA-protein interactions within germ granules.

## Acknowledgements

We thank members in the Tang lab for discussion and critical comments, M. Kearse for providing mammalian expression vectors and sharing equipment on cell culture, L.C. Tu for sharing U2OS cells, C.C. Mello and H.C. Lee for sharing *C. elegans* strains, OSU Neuroscience Imaging Core for instruments (S10OD010383 and S10OD026842), the Caenorhabditis Genetics Center for providing the *C. elegans* strains (P40OD010440). I. Price was supported by Center for RNA Biology graduate fellowship and presidential fellowship at The Ohio State University. This work was supported by the NIH Maximizing Investigators’ Research Award (R35 GM142580) and NSF (MCB 2420329) to W. Tang.

## Author contributions

I.F.P. and W.T. conceived and designed the experiments. I.F.P., C.W. and W.T. preformed the experiments. I.F.P., C.W. and W.T. generated Figures. I.F.P., and W.T. wrote and edited the manuscript.

## Supplementary Information

**Supplementary Fig. 1.** LOTUS-domain proteins and Vasa proteins. a. Schematics of LOTUS domain proteins and their domains in *C. elegans*, *D. melanogaster*, and *H. sapiens*. b. Schematics of Vasa and its orthologs *in C. elegans, D. melanogaster,* and *H. sapiens*. c. Fraction of endogenously tagged GFP::GLH-1 signal localizing to condensed foci in the germ lines of worms in the indicated mutant backgrounds. Number of germ lines quantified is indicated as n. Student’s t-test **** p-value ≤ 0.00005. d. Maximum intensity projection of a confocal Z stack showing the body wall muscle of a fixed worm ectopically expressing EGGD-1::GFP under the muscle-specific *myo-3* promoter. Scale bar = 20 μm. e. Single confocal slice of a nucleus outlined by the dashed box in (d). Scale bar = 5 μm. f. Maximum intensity projection of a confocal Z stack showing the body wall muscle of a fixed worm ectopically expressing GLH-1::mCherry under the muscle-specific *myo-3* promoter. Scale bar = 20 μm. g. Single confocal slice of a nucleus outlined by the dashed box in (f). Scale bar = 5 μm.

**Supplementary Fig. 2.** Quantification of relative EGGD-1 and GLH-1 expression levels. a. Plot of EGGD-1 and GLH-1 protein abundance, in arbitrary units, as determined by Tan et al. using mass spectrometry^29^. b. Plot of fluorescence signals in germ granules of strains expressing EGGD-1::GFP and GFP::GLH-1 levels. c. Quantitative Western blot to determine the relative abundance of EGGD-1 and GLH-1. The indicated number of gravid adult worms expressing EGGD-1::3xFLAG and 3xFLAG::GLH-1 were loaded into the corresponding wells. GLH-1 to EGGD-1 ratio was estimated by interpolation between the intensity of 1 and 2 3xFLAG::GLH-1 worms loaded.

**Supplementary Fig. 3.** Ectopic expression of wild-type and mutant GLH-1::mCherry. Maximum intensity projection of confocal slices spanning the heads of worms trangenically expressing EGGD-1::GFP with wild-type, T663A, or DQAD GLH-1::mCherry. Dashed boxes outline nuclei shown in Fig. 3d. Images are representative of at least 8 worms imaged.

**Supplementary Fig. 4.** Purification of eLOTUS and GLH-1 proteins. Coomassie blue staining of purified recombinant 6xHis::MBP::GLH-1, 6xHis::MBP::EGGD-1 eLOTUS domain, 6xHis::MBP:EGGD-1 eLOTUS (L88D).

**Supplementary Fig. 5.** Expression of EGGD-1::GFP and PGL-1::RFP. a. Pearson correlation coefficient of EGGD-1::mCardinal and PGL-1::GFP signal localized around EGGD-1 positive muscle nuclei. Each point indicates correlation coefficient of signal around a single nucleus. 5 nuclei were quantified per worm, n indicates the total number of nuclei analyzed. Student’s t-test, p-value****≤0.0005. b. Same as (A) but with PGL-3::GFP. c. Plot showing surface-to-rachis intensity ratio of EGGD-1::GFP in the indicated mutant and RNAi conditions. n indicates number of germ lines quantified. Student’s t-test, * p-value ≤ 0.05, ns: not significant. d. Images depicting single germline nuclei from live worms expressing endogenously tagged EGGD-1::GFP and PGL-1::RFP in the indicated mutant or RNAi conditions. Scale bar = 5μm. e. Pearson correlation coefficient of EGGD-1::GFP and PGL-1::tagRFP signal around individual germline nuclei. Each point indicates correlation coefficient of signal around a single nucleus. 5 nuclei were quantified per worm, n indicates the total number of nuclei analyzed. Student’s t-test, p-value < 0.00005.

**Supplementary Fig. 6.** Sequences and structures of DDX4 and TDRD5. a. Structural alignment between the Alphafold predicted the GLH-1 and DDX4 helicase domains. b. Sequence alignment between the EGGD-1 eLOTUS1 and TDRD5 eLOTUS domains. Mutated interfacial residues EGGD-1 L88 and TDRD5 I87 are indicated by the black arrow. c. Structural alignment between Alphafold prediction of EGGD-1 eLOTUS1 and TDRD5 eLOTUS domains. The C-terminal extended α5 helix is indicated and the predicted location of EGGD-1 L88 and TDRD5 I87 are indicated by the black arrow.

**Supplementary Table S1:** *C. elegans* Strains.

**Supplementary Table S2:** Oligo List.

## Materials and Methods

### Strains and maintenance

*C. elegans* strains were maintained on Nematode Growth Medium (NGM) under standard growth conditions at 20 °C^47^. Adult hermaphrodites were used for all experiments. A list of strains used in this study is located in Supplementary Table S1.

### RNAi by dsRNA feeding

NGM plates containing 50 μg/ml Ampicillin and 2.5 mM isopropyl β-D-1-thiogalactopyranoside (IPTG) were seeded with overnight cultures of HT115 carrying *glh-1* sequences cloned into the L4440 RNAi vector and kept at room temperature for 3 days to allow for the expression of dsRNA. 2-3 L4 worms were picked to the seeded plates and the F1 generation were used for downstream experiments.

### CRISPR genome editing

Point mutations at the endogenous loci of *eggd-1* and *glh-1* were generated using CRISPR-Cas9 mediated gene editing as described^48^. Briefly, guide RNA target sites adjacent to the mutation sites were assembled in to Cas9 RNP complexes and injected with synthetic single-stranded HDR templates (IDT) bearing the desired mutation in addition to the *rol-6* co-injection marker^49^. F1 worms were single picked to new plates and genotyped by PCR and restriction endonuclease digestion to identify mutant alleles. The list of oligos used for gene editing can be found in Supplementary Table S2.

### Plasmid construction

Plasmids used for transgenic expression of germ granule proteins in *C. elegans* muscle were generated using the PCFJ104 plasmid backbone containing mCherry flanked by the *myo-3* promoter and *unc-54* 3’UTR^50^. GFP, mCherry and mCardinal sequences were amplified using primers that introduced a 5’ XbaI site and 3’ SacI site in addition to restriction sites at the 5’ and 3’ end of the fluorescent protein CDS for cloning. The mCherry sequence in PCFJ104 was digested out using XbaI and SacI, and replaced with GFP, mCherry, or mCardinal. *eggd-1, glh-1, pgl-1,* and *pgl-3* cDNAs were cloned to these plasmids and that were subsequently used to generate transgenic strains. Primers to

introduce point mutations to *glh-1* and *eggd-1* cDNAs were designed using the NEBase Changer tool and the plasmid was mutagenized using the NEB Q5 site-directed mutagenesis kit. Mutations were confirmed by Sanger sequencing. EGGD-1 LOTUS1 cDNA, and GLH-1 helicase domain cDNA were cloned to the pET-His6-MBP-TEV-LIC cloning vector by ligation independent cloning for recombinant expression in bacteria. TDRD5 cDNA with GFP sequences and DDX4 cDNA with mCherry sequencs were cloned into the pCDNA3.1 vector. Oligos used for cloning and site directed mutagenesis can be found in Supplementary Table S2.

### Generation of transgenic worm strains

All transgenic strains were generated in *unc-119(ed3)* mutant worms. Transgenic strains were made by injecting *unc-119* worms with transgene plasmids mixed equally with PCFJ151 at 200 ng/µl. Injected worms were single picked to NGM plates and incubated at 20°C for 7-9 days, or until mobile F2 worms could be found on the plate. One mobile F2 worm per injection plate was picked to establish each transgenic transmission line.

### Confocal fluorescence microscopy

Slides with live worms were prepared by picking adult animals to a droplet of M9 buffer with 2.5 mM levamisole on 4% agarose pad. Worms were covered with 1.5H cover glass (THORLABS, CG15CH) and the slides were sealed using VALAP (1:1:1 vaseline:lanolin:paraffin). Fixed worm samples were prepared by suspending mixed-stage L4 and Adult worms with M9 + 0.05% Tween20 and transferring them to a 1.5 ml microcentrifuge tube. The worms were washed 3 times in M9 buffer to remove bacteria and placed in –20°C methanol for 20 minutes. Methanol was removed and PBS + 3.7% paraformaldehyde (PFA) was added. Worms were fixed in PFA solution for 30 minutes rotating at room temperature. The PFA solution was removed, and the worms were washed 3x in PBS before being resuspended in 50 μL of Vectashield antifade mounting medium with DAPI (Vector Laboratories, H-1200-10). The fixed worms were mounted to slides by pipetting a 3-5 μL to a slide and covering the droplet with 1.5H cover glass and sealing the slide with VALAP. Live imaging of *C. elegans* strains was performed using an inverted Nikon Ti2 microscope equipped with a CrestOptics X-Light V3 spinning disc confocal module and a Hamamatsu Orca Fusion C14440-20UP detector operated by NIS Elements AR using a Nikon CFI Plan Apo VC 60X WI objective. Whole fixed transgenic worms were imaged using the same setup with a Nikon CFI Plan Apo VC 60x Oil objective. Dissected gonads were imaged using Zeiss LSM 900 microscope equipped with a GaAsP detector operated by Zeiss Zen Blue 3.1 using a 63x Plan Apochromat oil immersion objective. For the preparation of mammalian cell slides, U2OS cells were collected from an existing culture and seeded directly onto coverslips in a 6-well plate. 40 hours after plasmid transfection, cells were washed with PBS, fixed with 3.7% (v/v) paraformaldehyde at room temperature for 15 minutes, and then washed again with PBS. Coverslips were mounted using Vectashield antifade mounting medium containing DAPI to stain the nuclei. Image processing was performed in ImageJ.

### Fluorescence recovery after photobleaching (FRAP)

FRAP was performed on live worms using a Zeiss LSM 900 microscope equipped with a GaAsP detector operated by Zeiss Zen Blue 3.1 using a 63x Plan Apochromat water immersion objective. ROIs were drawn around 2-5 foci of ectopically expressed EGGD-1 and GLH-1 per worm and bleached using the bleaching function in Zen and imaged every second for 100 seconds post-bleaching. FRAP in live U2OS transfected with TDRD5 and DDX4 plasmids was performed with a Nikon TiE microscope equipped with a Yokogawa CSU-W1 spinning disc confocal unit, an Andor iXon ULTRA 897BV EMCCD detector, and an Andor laser-galvo FRAPPA module operated by MetaMorph 7X Premier software using a Nikon 60x CFI Plan Apo VC water immersion objective. The cells were kept in BrainBits Hibernate B medium during imaging and maintained at 37°C using an Okolab Bold Line stage-top incubator. ROIs were drawn around TDRD5 foci and bleached using the targeted illumination tool in Metamorph. Cells were imaged every second for 120 seconds post bleaching. Intensity data were pulled from images using the measure function in ImageJ, analyzed using custom R scripts and plotted using the ggplot2 package in R.

### Fluoresence *in-situ* hybridization (FISH)

Whole-worm FISH was performed using mixed-stage L4 and adult worms. Worms were suspended from NGM plates with M9 + 0.05% Tween20, transferred to low-binding 1.5 ml microcentrifuge tubes and washed 3 times with M9 to remove bacteria. Worms were then washed once with ddH_2_O, suspended in –20⁰C methanol and kept at –20°C for 20 minutes. Methanol was removed by pipetting and 1 ml of PBS + 3.7% w/v PFA was added to the worm pellet. Tubes were wrapped in aluminum foil and rotated at room temperature for 45 minutes. After fixation, the PFA solution was removed by pipetting and the worms were washed 4 times in PBS, suspended in 75% ethanol, and incubated at 4°C for 4 hours. Worms were then washed once with Stellaris wash buffer A and incubated overnight at 37°C in Stellaris hybridization buffer with 1 μM of a 5’ Cy5-labeled DNA probe targeting the spliced-leader 1 consensus sequence^35^. The following day, hybridization buffer was removed, and worms were washed for 1 hour in Stellaris wash buffer A, followed by 1 hour in Stellaris wash buffer B. Wash buffer B was removed, and worms were suspended in 50 μL of Vectashield antifade mounting medium with DAPI and transferred to slides for imaging. FISH with dissected gonads was performed as described above with the following changes: After washing with M9, synchronized adults were dissected in M9 + Tween20 buffer with 0.5 mM levamisole before suspending in –20°C methanol. Fixation in PFA was reduced to 20 minutes.

### Western blotting

100 synchronized adult *3xflag::glh-1* or *eggd-1::3xflag* worms were suspended in M9 + 0.05% Tween20 and washed 3 times with M9 to remove bacteria. Worms were resuspended in 60 μL 50 mM Tris-HCl pH 7.5 and 20 μL of LDS buffer and 5 μL of 1M DTT were added. Samples were heated to 95 °C for 5 minutes and centrifuged at 20000xg before loading into a SDS PAGE gel. The protein was transferred to a PVDF membrane at 30V for 1 hour and blocked in PBSt + 3% milk (PBS + 0.1% Tween20) at room temperature for one hour. anti-FLAG M2 (Sigma F3165) antibody was used at a dilution of 1:2000 and goat anti-mouse secondary (Abcam ab97023) was used at a concentration of 1:10000.

### Purification of recombinant proteins

6xHis-MBP-tagged EGGD-1 LOTUS domain and GLH-1 helicase domain were obtained by recombinant expression in *E. coli* and purified by batch dual affinity. Overnight cultures of Rosetta2 (DE3) cells carrying EGGD-1 or GLH-1 expression vectors were used to inoculate LB broth supplemented with 50μg/ml kanamycin and 34 μg/ml chloramphenicol and grown at 37°C to an OD of 0.6. Upon reaching an OD of 0.6, the cultures were transferred to an ice bath for 30 minutes after which Isopropyl β-d-1-thiogalactopyranoside (IPTG) was added to a final concentration of 0.5mM. Cells were incubated overnight at 18°C and harvested the following day by centrifugation. Bacterial pellets were suspended in lysis buffer (300mM KCl, 50 mM Tris-HCl pH 7.5, 10 mM imidazole, 0.5mg/ml lysozyme) and lysed by sonication. Lysates were centrifuged in a Beckman Coulter JA-20 rotor at 18000 RPM for 30 minutes at 4°C. Centrifuged lysate was applied to 1 ml of Ni-NTA resin in a 50 ml conical tube and rotated at 4°C for 10 minutes. Tubes were centrifuged at 1000xg to pellet the resin and supernatant was removed. The resin was washed twice for 5 minutes in 45 ml of wash buffer (300mM KCl, 50 mM Tris-HCl pH 7.5, 20 mM imidazole). After the second wash, the total volume was applied to a BioRad EconoPac gravity flow column. Once the wash buffer had passed through the protein was eluted from the resin in 5ml of elution buffer (300mM KCl, 50 mM Tris-HCl pH 7.5, 250 mM imidazole). Eluate was diluted 10-fold in MBP buffer (300mM KCl, 20mM HEPES-KOH pH 7.4, 1mM EDTA), applied to 1 ml of NEB amylose resin (E8021S) in a 50 ml conical tube and rotated at 4°C for 30 minutes. The resin was washed twice in 45 ml of MBP buffer and transferred to a gravity flow column. Protein was eluted from the resin in 5 ml MBP buffer supplemented with 10 mM maltose. Eluate was passed through a 0.2 μm filter and concentrated using Amicon Ultra 30k centrifugal filters (Sigma-Aldrich, UFC803008). Protein was mixed with 1/10 volumes of 50% glycerol, aliquoted and flash-frozen in liquid nitrogen before storing at –80°C.

### Malachite green ATPase assay

1 μM EGGD-1 LOTUS domain and 5 μM GLH-1 helicase domain were incubated at 25°C in assay buffer (60mM KCl, 50 mM NaCl, 25 mM HEPES, 5 mM MgCl_2_, 1 mM DTT, 100 μg/ml Bovine Serum Albumin) supplemented with 1 mM ATP and 20 ng/μL of a 200 nt *in-vitro* transcribed RNA substrate. At each time point, the ATPase reaction was terminated by transferring 10 μL of the reaction to 160 μL of 10 mM EDTA pH 7. The presence of free phosphate in the terminated reactions was assayed using a malachite green phosphate assay kit according to the manufacturer’s instructions (Sigma Aldrich, MAK307). In brief, 80 μL of terminated reaction was mixed with 20 μL of malachite green working reagent in an optically clear 96 well plate and incubated at room temperature for 30 minutes for color to develop. Absorbance in wells was measured using a Promega GloMax Discover microplate reader and data were plotted using ggplot2 package in R.

### Electrophoretic mobility shift assay (EMSA)

EMSA assays were conducted using 1 μM GLH-1 helicase domain and the indicated concentrations of EGGD-1 LOTUS domain in assay buffer supplemented with 10 mM AMP-PNP and 100 nM 5’-Fluorescein labeled 30 nt poly-U RNA substrate. Reactions were assembled on ice and incubated at 20°C for 20 minutes. After incubation, reactions were placed back on ice and loaded into an 8% native PAGE gel made with 0.5x tris-borate-magnesium buffer (45 mM Tris HCl, pH8.3, 45 mM boric acid, 2.5 mM MgCl_2_) and run at 100 V for 60 minutes at 4°C. Gels were imaged using an Azure Sapphire Biomolecular Imager. The intensity the bands was quantified using ImageJ and the data were plotted using the ggplot2 package in R.

### Cell culture and transfection

U2OS cells were maintained in high-glucose Dulbecco’s Modified Eagle’s Medium (Sigma-Aldrich, D6429) supplemented with 10% heat-inactivated fetal bovine serum (Sigma-Aldrich, 12306C) and 10,000 U/ml penicillin-streptomycin (Gibco, 15140122) at 37 °C with 5% CO₂. For transient transfection, U2OS cells were seeded into a 6-well plate one day prior to transfection and transfected with 2 μg of TDRD5::GFP and/or DDX4::mCherry expression plasmids using ViaFect (Promega, E4981) according to the manufacturer’s instructions.

### Image analysis

#### Quantification of condensed GLH-1 fraction

A section of the germline encompassing mid-pachytene germ cells was cropped from z-stacks encompassing the surface to halfway through the gonad for each strain analyzed. The signal quality from the top half of the gonad is much higher than the bottom half of the z-stack. For this reason, symmetry was assumed and only the top half of the stack was used to quantify the condensed protein fraction. Images of GFP::GLH-1; *eggd-1* mutant worms were used to determine the threshold used for segmentation of condensed foci for all strains. To determine the threshold value, a maximum intensity projection was performed for each image and the auto-threshold value was determined using the “Li” threshold algorithm. The median threshold value was applied to each image using the “3D Simple Segmentation” plugin in ImageJ to generate a mask around condensed GLH-1 foci. The mask was used to generate an image that only contained the condensed signal, and a sum intensity projection was applied to the original z-stack and the z-stack of the condensed signal. The total intensity of each image was measured, and the quotient of the condensed/total signal was used to plot the condensed GLH-1 fraction.

#### Quantification of surface-to-rachis signal intensity

One z slice at an optical cross section of the germline was selected per worm analyzed. An ROI was drawn tightly around the dorsal row of germ cell nuclei and the average signal intensity was measured to determine the average intensity at the surface of the germ line. The average intensity of a section of the same length was measured in the germline rachis. The ratio was determined by dividing the average intensity at the surface by the average intensity in the rachis and plotted using the ggplot package in R.

#### EGGD-1 and PGL-1/PGL-3 colocalization

Ectopically and endogenously expressed EGGD-1 and PGL-1/PGL-3 colocalization were quantified using the same approach. ROIs were drawn around individual nuclei and used to generate a new image. The image channels were split, and a maximum intensity projection was performed with each channel. The maximum intensity projection was used to determine an auto-threshold value for the channel using the “Li” threshold algorithm. This threshold value was applied to the full z-stack to generate a mask using the “3D Simple Segmentation” tool. The masks of both channels were added together to make a combined mask that was used for colocalization analysis using the Coloc2 plugin.

